# Combined pharmacotherapy suppress ferroptosis and inflammation after neonatal intraventricular hemorrhage

**DOI:** 10.64898/2026.11.09.765451

**Authors:** Michelle Sunshine, Vanessa Castro Diaz, Christina LaGamma, Furong Hu, Sansar Sharma, Selvakumar Subbian, Libor Velisek, Jana Veliskova, Ravi Sachidanandam, Govindaiah Vinukonda, Edmund F. La Gamma

## Abstract

Neonatal intraventricular hemorrhage (IVH) is a common complication of extremely premature newborns, and results in reduced neurogenesis, white matter injury, cortical atrophy, and subsequent cerebral palsy, learning and memory disabilities. However, the impact of IVH on the hippocampus, an area critical for memory and learning capabilities throughout life, remains relatively unexplored. Using a preterm rabbit pup model that structurally resembles human perinatal brain development, we report, widespread hematoma and iron accumulation in the hippocampal dentate gyrus (DG) following IVH. This accumulation is facilitated by the anatomical positioning of the hippocampus within the unidirectional CSF flow pathway and the vulnerability of the immature dentate gyrus vasculature. Sulforaphane (SFN; an activator of nuclear factor erythroid 2 related factor 2 [Nrf2] transcriptional factor and anti-inflammatory pathways) and deferoxamine (DFN; an iron chelator) reduced iron deposition, free hemoglobin, and activated glial cells (sp. astrocytes and microglia), particularly in the DG molecular layer and hilar regions. The attenuated inflammatory responses correlate with decreased ferroptosis-mediated neuronal death, and substantial restoration of the dysregulated gene pathways associated with inflammation, immune activation, and neuronal cell death in the developing hippocampal/DG region. This was associated with a gene profile favoring ferroptotic cell death rather than apoptosis or autophagy pathways. Moreover, we also identified a regional specificity to certain effects as no astrogliosis was observed in hippocampal CA1 to CA4 regions after IVH or when compared with healthy controls. Our study is the first to demonstrate hippocampal hematoma and iron accumulation associated pathologic mechanisms following neonatal IVH. The results provide mechanistic insights into hippocampal injury as well as a therapeutic proof-of-concept for targeting iron toxicity and inflammation in the developing brain to mitigate injury. These results justify extending analyses of injury reversal to long-term postnatal ages and behavior studies to assess therapeutic potential. The study illustrates novel therapeutic interventions that attenuate early mediators of injury that improves recovery of hippocampal/DG formation/maturation and neuroprotection after neonatal IVH. We speculate that these protective effects may prolong donor stem cell survival thus, potentiating benefits of living cellular therapy in the future.

## 1. INTRODUCTION

Intraventricular hemorrhage (IVH) is a common complication of extremely premature newborns, and results in reduced neurogenesis, white matter injury (WMI), cortical atrophy, and subsequent cerebral palsy, learning and memory disabilities with increased long-term health care costs in surviving infants.^1–5^ Despite many advances in neonatal and perinatal care toward prevention of IVH, there are no effective interventional options to recover normal brain development in humans.

IVH commonly arises from the germinal matrix, a highly cellular, developmentally transient structure, where all new precursor cells are born, proliferate, and migrate during fetal and perinatal brain development. ^6, 7^ IVH results from the fragility of immature vasculature with increasing susceptibility to various detrimental perinatal events^1^. Specifically, immature parenchymal support structure and autoregulatory control of cerebral blood flow contribute to pressure-dependent blood vessel rupture that leads to extravasation of red blood cells (RBC) into the brain parenchyma and subsequently, into the glymphatic^8^ and ventricular CSF drainage systems. Later, RBC lysis releases free iron, excess heme, and globin proteins as toxic compounds that induce widespread collateral inflammatory damage. Most importantly, recovery from the injury process interrupts the normal progression of cell lineage differentiation, synapse formation and structural integrity of the developing brain. In the past, the majority of studies of IVH focused on WMI and hydrocephalus, however, the hippocampus is a periventricular cortical gray matter structure important for memory formation and learning capabilities, yet, it has received comparatively less attention. ^9^ After IVH, unidirectional flow of CSF and glymphatic drainage,^8^ facilitates the clearance of hematoma breakdown products, but can injure the hippocampus in tandem due to its periventricular location and after rupture of immature hippocampal blood vessels. Since the morphogenesis of the hippocampal-dentate gyrus (DG) peaks during the antenatal third trimester; ^10^ early exposure to adverse conditions in this region causes permanent detrimental changes in its subsequent development.

There are very few studies in the hippocampus characterizing hemorrhagic hematoma, iron accumulation, cell death mechanisms, or astrocytes and microglia mediated inflammation after spontaneous IVH. In a rat model using exogenous hemoglobin injection (to mimic intraventricular hemorrhage), a study identified neuronal cell death, increased c-Jun N-terminal kinase (JNK) activation and co-localized hemoglobin receptor CD163 leading to neurodegeneration of the Cornu Ammonis area-1 (CA-1) neurons. ^11, 12^ In another report, the authors injected whole blood into the hippocampus of postnatal day 4 (P4) rat pups, and observed increased neuronal cell death and activation of BDNF-TrkB-CREB signaling.^13^ Additional studies performed transcriptome and microRNA analyses on isolated microglia and neurons revealing time-specific cascades for appearance of neuronal-specific miRNA including those relevant to development of the CA-1 region.^14–16^ Treatment interventions focused on interruption of these detrimental consequences are not well studied.

In the present report, we evaluated the effects of combined pharmacological treatment with deferoxamine (DFN, iron chelator) and sulforaphane (SFN), an activator of Nrf2 transcription factor (induces anti-inflammatory cytokines) on the pattern of injury in the hippocampus. Sulforaphane modifies Keap1, freeing Nrf2 to enter the nucleus and bind to antioxidant response elements (ARE) that induce phase II detoxification enzymes such as hemeoxygenase-1 (HMO-1).^17^ We hypothesized that eliminating injurious toxicants and augmentation of anti-inflammatory pathways (sp. ARE responsive genes) would attenuate free radical iron toxicity, reduce proinflammatory collateral injury, attenuate cell death, and therefore, help restore normal hippocampal development. We found that drug treatment reduced: free iron levels, ferroptotic cell death, reactive astrocytes, and microglial infiltration, and altered gene expression profiles. These changes indicate that combinatory DFN and SFN treatment promotes a less toxic microcellular environment. We speculate that early pharmacological removal of hemorrhagic toxicants may enhance survival of stem cells used as “*living cell therapy*” in a changing microenvironment, thereby preserving their adaptive capacity as anti-inflammatory agents promoting recovery of normal brain growth.

## 2. METHODS

### 2.1 Premature rabbit model of Glycerol-induced GMH-IVH

All rabbit experiments and treatment protocols were approved by New York Medical College Institutional Animal Care and Use Committee (IACUC). Timed pregnant New Zealand white rabbits were purchased from Charles River Laboratories Inc. (Wilmington, MA, USA) and E29 premature rabbit pups were delivered by Cesarean-section and pups were maintained and fed according to our previously published protocols.^3^ At 4-6 hours of postnatal age, newborn rabbit pups were injected with 50% intra-peritoneal glycerol (6.5 g/kg), which induced IVH. Head ultrasound was performed at 24h postnatal age to determine the presence and severity of IVH (Ultra-sonographic Imaging System, Zonare Medical System. CA, USA).^7, 18^ Rabbit pups were assigned to one of 3 experimental groups: Control (no glycerol injected healthy controls and glycerol injected with no IVH pups equally), Glycerol-IVH + saline injected pups and Glycerol- IVH + sulforaphane (SFN)-deferoxamine (DFN); see figure legends^19^.

### 2.2 Tissue collection and processing

As in our previous publications,^3, 7, 18^ the total hippocampal thick tissue slice (about 1 cm) was dissected under light microscope using a brain matrix slicer (Cat # BSRAS001-1, ZIVIC Instruments, Pittsburg, PA, USA) and coronal sections were cut for immunohistochemistry (IHC), immunofluorescence (IF) studies, and for RNA and tissue lysates.^3, 7, 18^ Cerebrospinal fluid (CSF) and plasma samples were collected prior to tissue dissection. Samples were snap-frozen on dry ice and stored in -80° C until analysis.

### 2.3 Histological evaluation of hematoma and iron accumulation

Hematoxylin and eosin staining (H&E staining) and tissue iron staining (Abcam kit; cat # ab150674) were performed using fixed coronal sections. ^3, 7, 18^. We evaluated every 10th section from the hippocampus for a minimum of 8 serial sections from anterior to posterior per sample in each group.

### 2.4 Quantification of free hemoglobin and iron deposition

Hippocampal tissue lysates and CSF collected were analyzed for free hemoglobin (Hb) (Cat # ab272533, Abcam kit, USA) and iron (Cat # ab83366, Abcam kit, USA). Total free hemoglobin and free iron were calculated as described in the kit’s protocol.

### 2.5 Immunohistochemical and Immunofluorescence staining

IHC and IF staining were performed ^3, 7, 18^ using the following antibodies for Neuronal Nuclei (NeuN) staining (Cat # ab104224, Abcam), Iba-1 (Cat # ab5076, Abcam) to assess immune cell infiltration by microglia, and GFAP (Cat # G3893, Sigma Aldrich) to assess astrogliosis. The sections were counter stained with DAPI nuclear stain. TUNEL staining (Cat # 17-141, EMD Millipore/Sigma, St. Louis, MO, USA) was used to assess for neuronal cell death when merged with NeuN staining.

### 2.6 Cell density and quantification procedures

We performed all cell counting on fixed coronal sections as described previously. ^3, 7, 20^ We used 20X/40X objectives for images and counted 2-3 alternate sections covering all 3 layers of the DG for each pup. Cell counts were performed independently by two investigators (blinded to the experimental groups) using Image-J software with a grid. We counted 4 sections in each pup and 5 pups from each group. Data are presented as mean cell count (mean ± s.e.m) for TUNEL, NeuN with TUNEL, IBA-1 and GFAP.

### 2.7 RNA Isolation and TaqMan gene expression assay

Dissected hippocampal/DG tissue total RNA extraction was performed using Qiagen RNeasy Mini Kit (cat #74104) procedure following our published methods.^19^ TaqMan gene expression was performed using ABI quanta studio-3 (Thermo Fisher Scientific, Waltham, USA) as previously described and published method. ^3, 20^ The following single tube TaqMan probe-primer mixes for each target were used: *Glyceraldehyde-3-phosphate dehydrogenase (GAPDH) ID: Oc03823402_g1; Glial fibrillary acidic protein (GFAP) ID:OC03823402-G1; Hemeoxygenase-1 (HMOX1) ID: Oc03396031_m1; Transferrin (TF) ID: Oc03399304_m1; Transferrin receptor (TFRC) ID: Oc06680198_m1; Acyl-CoA synthetase long chain family member 4 (ACSL4) ID: Oc06767952_m1; Nuclear receptor coactivator 4 (NCOA4) ID: Oc06781088_m1, STEAP3 metalloreductase, ID: Oc06729595_m1; Lysophosphatidylcholine acyltransferase 3 (LPCAT3) ID: Oc06732656_g1; Iron responsive element binding protein 2 (IREB2) ID: Oc06677954_m1; Prostaglandin-endoperoxide synthase 2 (PTGS2) ID: Oc03398292_g1; BCL2 related protein A1 (BCL2A1), ID: Oc06717379_m1; Solute carrier family 7 member 11 (SLC7A11), ID: Oc06683681_m1).* All probe-primers were purchased from Thermo-Fisher Scientific, Waltham, MA, USA. Data analysis was completed using the efficiently corrected delta-delta CT method and the data was presented in percentages.

### 2.8 mRNA sequencing Analysis

#### RNA-Seq and pathway analysis

Rabbit hippocampal/DG total tissue samples were harvested from 1 cm thick coronal sections on postnatal days 3 and 7 starting from the anterior end to the posterior end and immediately snap frozen in liquid nitrogen and stored it at -80^0^C. Total RNA extraction was performed using Qiagen total RNA isolation kit following our published methods.^19, 3^ RNA-sequencing was performed using Ambion ERCC (External RNA Control Consortium) spike-in at the earliest stage (i.e. right after tissue collection) for internal control. The ERCC spike-in was used to identify the lower limit of detection and for normalization between samples in data analysis. We performed RNA-seq analysis using Illumina TruSeq stranded mRNA library prep kit for RNA-seq library preparation and used Illumina NextSeq 2000 system for sequencing at our Genomics Core Lab at New York Medical College. Gene expression and pathway analysis assessed after RNA-seq raw data was analyzed for identifying individual genes, differential expression and pathways and network analyses to reduce the complexity and identify differentially active pathways and networks. The hippocampus dissected mRNA sequence raw-data was deposited to GEO-NCBI (GSE 305615).

### 2.9 Statistical analyses

Scatter plot with bar graphs represent mean ± SD as described in the figure legends. Standard deviation (SD) was used to assess variation between experimental groups. Multiple groups were compared at two postnatal ages to evaluate the main effects of treatment and impact of age by one-way ANOVA using GraphPad Prism 6 (GraphPad Software, CA, USA). *Post-hoc* comparisons were done using Tukey’s test. A p-value of p < 0.05 was considered statistically significant.

## 3. RESULTS

### 3.1 Hematoma/free-hemoglobin accumulation during postnatal hippocampal/DG formation after IVH in premature rabbit pups

To assess whether IVH led to an accumulation of hemorrhagic bleed/hematoma deposition in the hippocampal dentate region, we examined forebrain coronal sections from the developing premature rabbit brain with and without hemorrhagic injury. ^18, 21, 22^ We used every 10^th^ section and assessed for hematoma accumulation using H&E stained sections. We observed normal developmental morphological appearance of hippocampal/DG sub-region formation in healthy, no IVH control pups at postnatal day 3 (Figure1A-B). In contrast, in IVH pups, there was extensive hematoma accumulation along with structural and cellular damage in the hippocampus/DG sub-regions at postnatal day 3 (Figure1C-D). Similar findings were observed at postnatal day 7 (Supplementary Figure-1). These results indicate significant hematoma debris during critical postnatal hippocampal formation.

**FIGURE 1:**
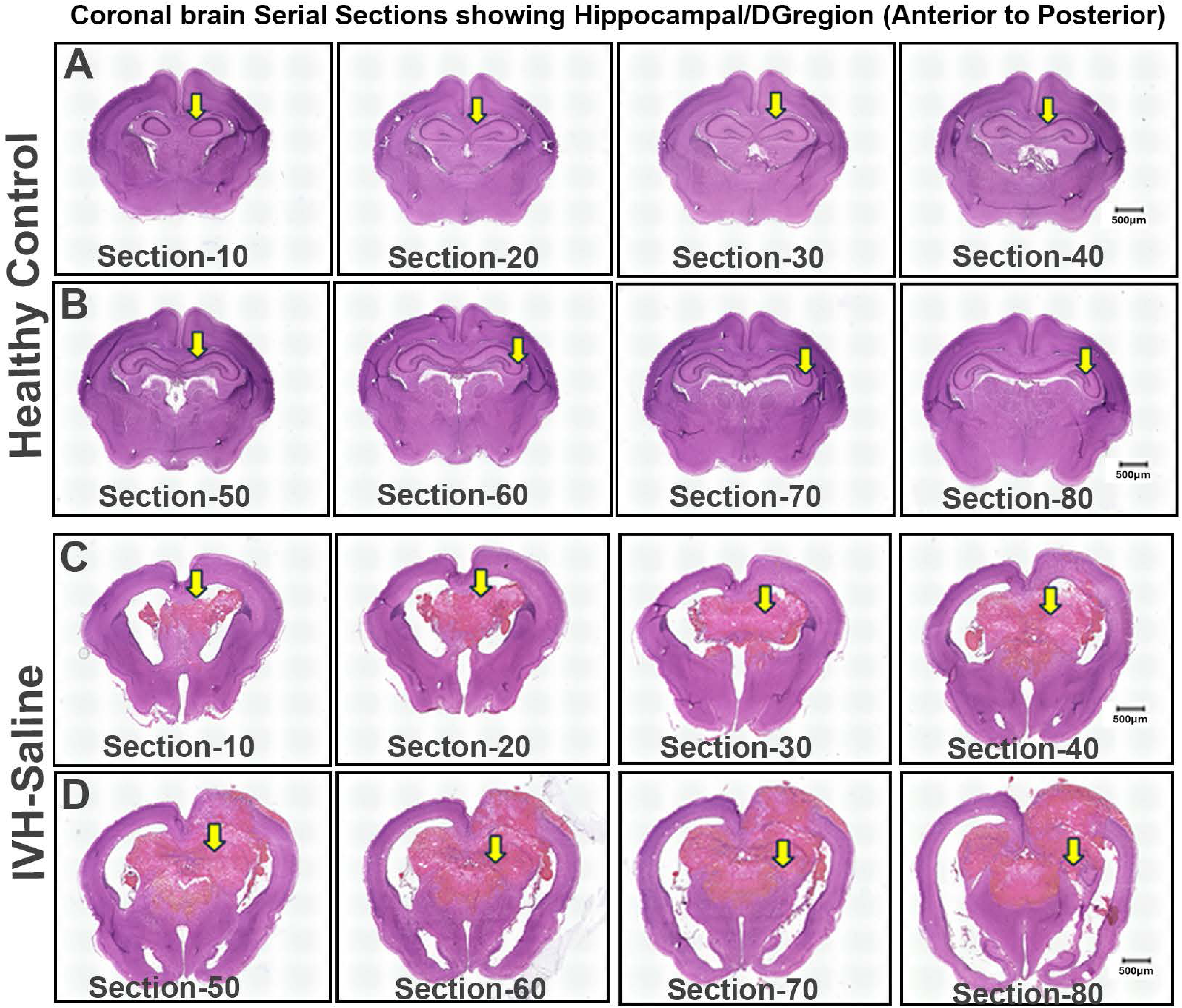
Postnatal hippocampal/DG formation and hematoma/debris accumulation after intraventricular hemorrhage in premature rabbit pups. A-B) Representative hematoxylin and eosin staining (H&E) on coronal serial section images showing hippocampal/DG sub-regions in healthy control premature rabbit pups without hemorrhage at postnatal day 3. C-D) Representative H&E staining on coronal serial section images showing huge hematoma accumulation in the hippocampal/DG regions in premature rabbit pups with morphologically damaged DG layers at day 3 Every 10th section was taken from anterior to posterior region for H&E stained sections and stitched image were taken at low magnification using KEYENCE microscope (Keyence corporation of America, Illinois, USA). The scale bar for all images 500 µm for stitched images.

### 3.2 Iron accumulation during postnatal/hippocampal/DG formation after IVH in premature rabbit pups

To assess whether IVH triggers an accumulation of iron in the hippocampus/DG region, we investigated hippocampal coronal sections with and without hemorrhagic injury. We observed iron deposition in the hippocampal/DG sub-region in IVH pups (Figure 2A-B vs 2E-F left hemisphere and 2C-D vs 2G-H right hemisphere) at postnatal day 3 compared with a lack of iron deposition in no IVH controls. After IVH, the hippocampi showed hematoma and iron accumulation at both postnatal days 3 and 7.

**FIGURE 2:**
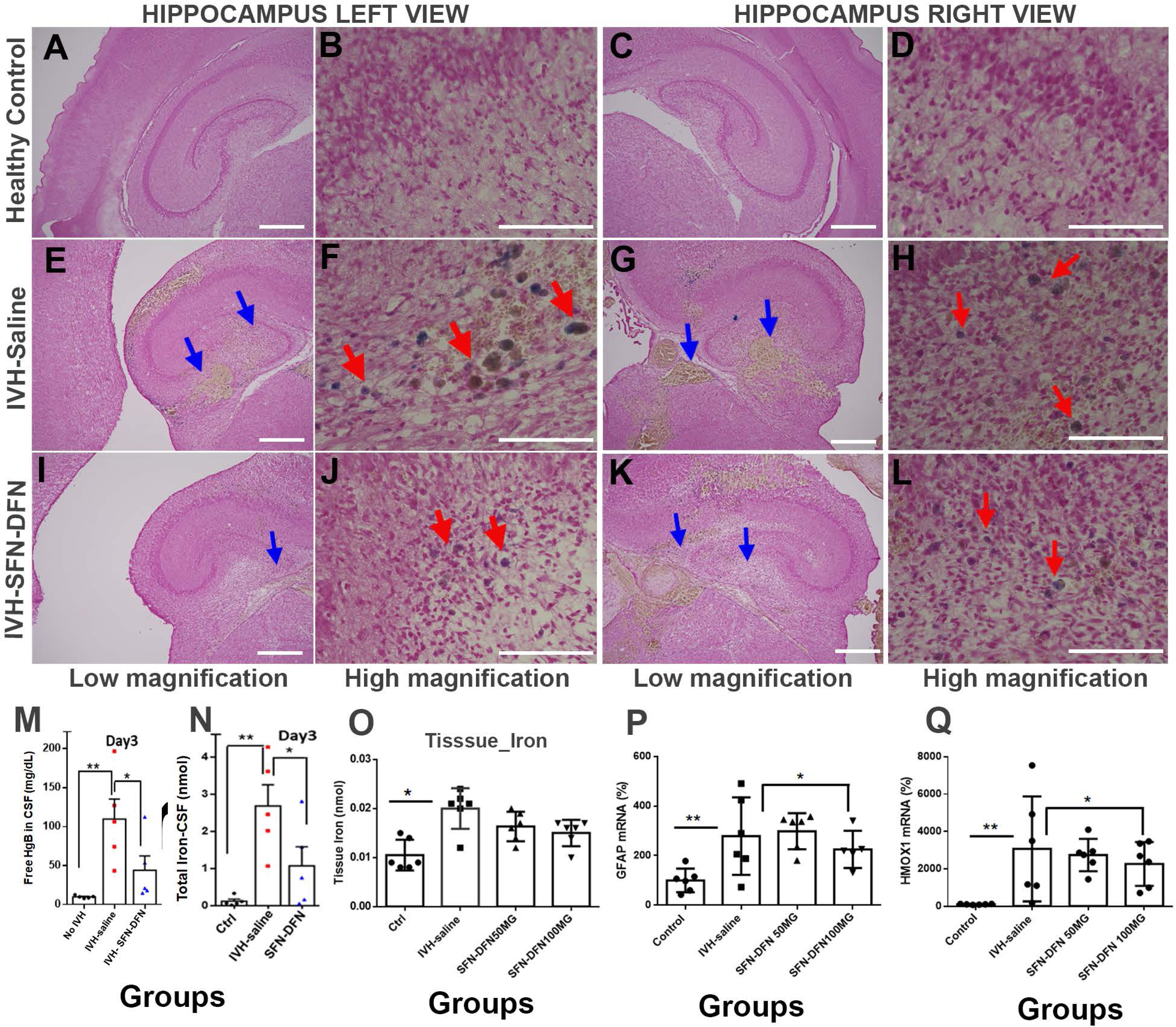
Sulforaphane (SFN) and Deferoxamine (DFN) treatment diminished IVH injury from iron accumulation during postnatal development of hippocampal/DG formation in premature rabbit pups. A-D) Representative iron staining images using H&E on coronal sections. Images show no iron deposition in hippocampal/DG sub-regions in healthy premature rabbit pups at postnatal day 3. The low and high magnification images are of left (A-B) and right (C-D) hippocampal hemispheres of the forebrain region in healthy premature pups at postnatal day 3. E-H) Representative iron staining images using H&E on coronal section. Images show iron deposition in hippocampal/DG sub-regions in premature rabbit pups with IVH at postnatal day 3. The low and high magnification images are of left (E-F) and right (G-H) hippocampal hemispheres of the forebrain region in premature pups with IVH at postnatal day 3. I-L) Representative iron staining images on coronal sections using H&E, images showing iron clearance after SFN-DFN treatment in hippocampal/DG sub-regions in premature rabbit pups with IVH at postnatal day 3. The low and high magnification images are of left (I-J) and right (G-H) hippocampal hemispheres of the forebrain region in premature pups with IVH at postnatal day 3. (Total tissue iron was assessed according to methods described in Methods (Cat # ab83366, Abcam, USA). The H&E stained images were created using a KEYENCE microscope (Keyence corporation of America, Illinois, USA). The scale bar illustrates low magnification images at 50 µm and high magnification images 25µm. The blue and red arrows indicate iron deposition in black/brown color. M-O) Scattered plot with bar graph showing cerebrospinal fluid, dissected hippocampal tissue lysates for free hemoglobin, and iron accumulation and, clearance after SFN-DFN treatment in premature rabbit pups at postnatal day 3. We used Abcam free hemoglobin (fHb) kits (Cat # ab272533) and iron assay kit (Cat # ab83366, Abcam, USA) to assess fHb and iron. The total fHb and iron was significantly increased in IVH compared with healthy controls in CSF, whereas after SFN-DFN treatment, they were both significantly reduced (M, N). In dissected hippocampal tissue lysates, we observed a significant increase in iron after IVH compared with controls and there was smaller change after SFN-DFN treatments, but it did not achieve significance (O). The data presented for CSF is mg/dL and in tissue lysate as nmol with mean ± S.D. Sample size of 5-6 in each group; *P < 0.05 and **P < 0.01. P-Q) Scatter plot bar graphs show TaqMan gene expression quantification using hippocampal/DG dissected isolated RNA in premature rabbit pups at day 3. The mRNA expression for GFAP (P) and HMOX1 (Q) significantly increased (p < 0.01) after IVH and this was significantly reversed for both, after 100 mg SFN-DFN treatment (D&E) at postnatal day 3. Sample size 5 in each group, P < 0.01 for CTRL vs IVH and P < 0.05 for IVH vs treatment for both GFAP and HMOX1 expression respectively at day 3. **SFN-DFN treatments:** Rabbit pups in the SFN-DFN treatment group were given SFN 25 mg/kg/dose intramuscular (IM) every other day along with DFN 50 mg/kg/dose subcutaneously (subQ) twice a day. In addition, a second set of experiments used a higher dose of DFN 100 mg/kg/dose (subQ) along with SFN 25 mg/kg/dose (IM) using a 1cc insulin syringe with 29-gauge needle. The pharmacological agents SFN purchased from EMD Millipore (Cat # 57421) and the iron chelator, DFN procured from Hospira, Inc (Cat # NDC0409-2336-01). The pups were sacrificed on postnatal days 3 and 7 experimental endpoints.

### 3.3 SFN-DFN treatment results in reduced hematoma and iron accumulation in the hippocampal/DG region after IVH

Excessive endogenous accumulated iron in cells and extracellular spaces is highly reactive and toxic. On postnatal day 3, coronal sections showed reduced iron accumulation after SFN-DFN treatment compared with IVH pups without treatment (Figure 2E-F vs I-J left and G-H vs K-L right). Since lysis toxicants appear in the circulating CSF and in the dissected hippocampal tissue lysate, we examined whether this would change after SFN-DFN treatment. We found increased free CSF hemoglobin levels in IVH pups without treatments, whereas after SFN-DFN treatment, we observed a significant reduction of free hemoglobin at postnatal day 3 (n= 4-5 samples in each group, p<0.01 between control vs IVH and p< 0.05 IVH vs SFN-DFN treated; Figure 2M). Quantification of total iron accumulation using CSF and tissues lysate indicated significant increases after IVH compared with no IVH controls for both CSF and tissue lysates (Figure 2N, O; n= 5 for CSF and n=6 for tissue samples in each group p< 0.01 for control vs IVH) at postnatal day 3. After SFN-DFN treatment, iron decreased significantly in CSF, (p< 0.05 for IVH vs SFN-DFN treated, n=5), although there was a decrease in levels of iron in tissue lysates, it did not reach control levels (Figure 2 O; n=6 samples in each group p< 0.05 for control vs IVH) at postnatal day 3.

### 3.4 SFN-DFN treatment results in reduced glial activation in the hippocampal/DG sub-region after IVH

After IVH, cell-free hemoglobin and iron are released and activate oxidative toxicity and inflammatory cascades resulting in reduced CNS cell proliferation and neuroglia cell death. Astrocytes and microglia respond to injury in the CNS by changing shape, enlarging and multiplying in a process referred to as astrocytosis/reactive gliosis with a shift toward inflammatory microglia (innate immune cells). Histologically, astrocytes (GFAP positive cells) and microglia (IBA-1 positive cells) were both increased (Figure 3A vs 3D panels) after IVH as compared with healthy controls at day 3. By comparison, after SFN-DFN treatment, IVH pups showed significantly reduced immune reactivity histologically and quantitatively for both reactive astrocytes and microglia (Figure 3D vs 3G panels). Further TaqMan gene expression assessment confirmed that GFAP mRNA was significantly higher in IVH compared with healthy controls and reduced significantly after SFN-DFN treatment at day 3 (Figure 2P, n=6 samples in each group p< 0.01 for control vs IVH; p<0.05 for IVH vs SFN-DFN higher dose).

**FIGURE 3:**
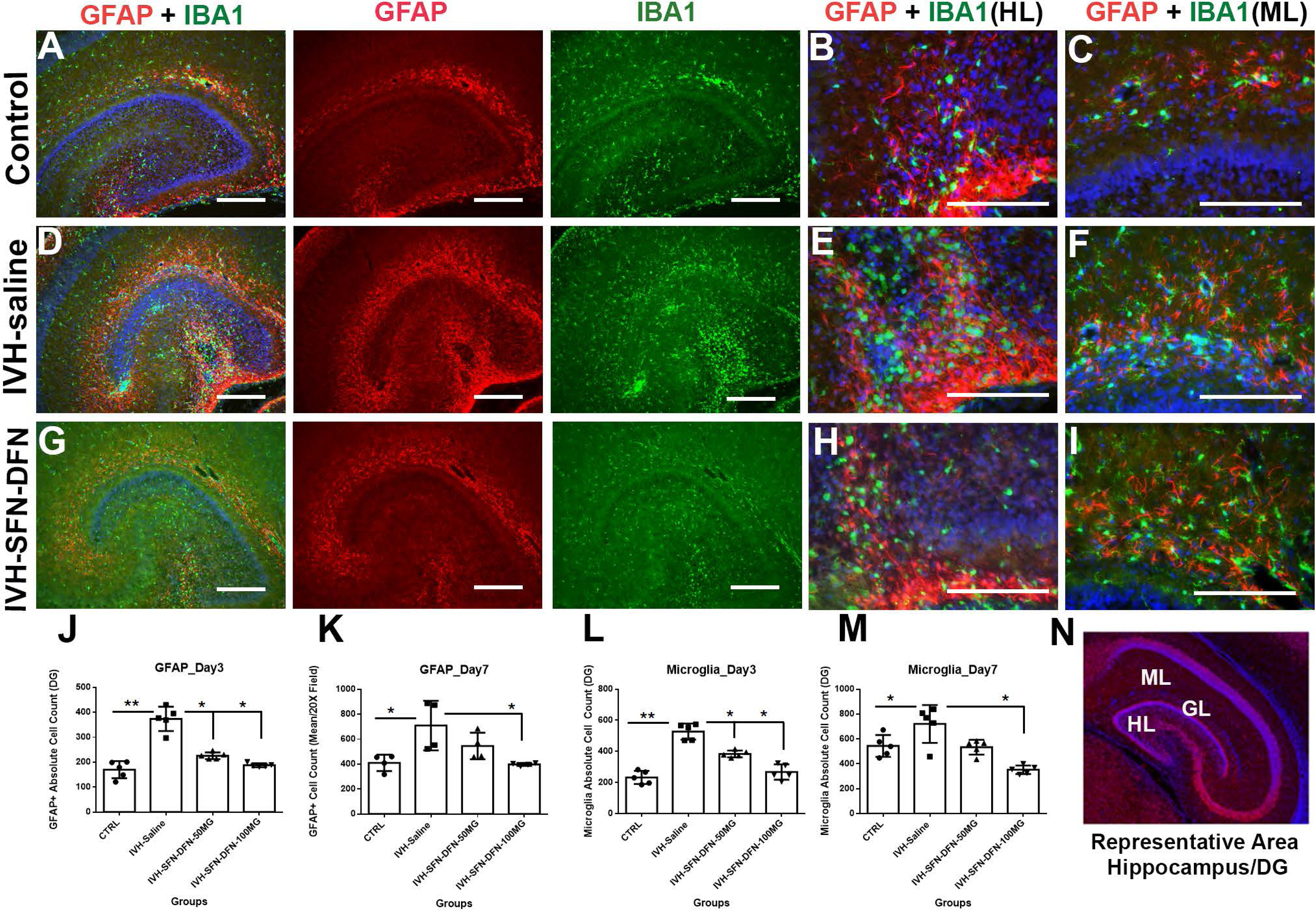
Sulforaphane and Deferoxamine treatment suppressed glial cell activation (astrocytes, microglia) after IVH injury during postnatal development of the hippocampal/DG region in premature rabbit pups. A-C) Representative immunofluorescence-stained images on coronal sections using GFAP (astrocyte marker) and IBA-1 (microglia marker) hippocampal/DG sub-regions in healthy control pups at postnatal day 3. Normal distribution of astrocytes and microglial immunoreactivity was observed in the hippocampal hemisphere (A), hilus and molecular layers (B, C) of the DG sub region in the healthy controls. D-F) Representative immunofluorescence-stained images on coronal sections using GFAP (astrocyte marker) and IBA-1 (microglia marker) hippocampal/DG sub-regions among IVH pups at postnatal day 3. Enhanced distribution of astrocytes and microglial immunoreactivity was observed in the hippocampal hemisphere (D), hilus and molecular layers (E, F) of the DG sub-region among premature pups with IVH. G-I) Representative immunofluorescence-stained images on coronal sections using GFAP (astrocyte marker) and IBA-1 (microglia marker) hippocampal/DG sub-regions among IVH pups treated with SFN-DFN at postnatal day 3. Suppressed distribution of astrocytes and microglial immunoreactivity was observed in hippocampal hemisphere for both merged and single stain images (G), hilus and molecular layers (H, I) of the DG sub-region among premature IVH pups treated with SFN-DFN. The immunofluorescence images were acquired using KEYENCE microscope (Keyence corporation of America, Illinois, USA). At postnatal day 3, 20 uM coronal sections were used for staining and the scale bars for all images was 250 uM. The low power magnification (triple labeled) merged images and single stain for GFAP or IBA-1 images were of the complete hippocampal hemisphere. High magnification objectives for triple labeled merged images are shown for hilus and molecular layers. A sample size of 5 pups were assessed in each experimental group. J-M) Scatter plot with bar graph shows absolute cell density quantification of GFAP positive astrocytes and IBA-1 positive microglia in the hippocampal/DG sub-region among 4 experimental groups of premature pups at postnatal days 3 and 7. The cell count was performed on coronal section images taken at 20X magnification in both hippocampal hemispheres. Sample size of 5 in each group, *p < 0.05,**p < 0.01. N) Representative immunofluorescence-stained images of coronal section using NeuN antibody (neurons) and IDAPI (nuclear marker) to show of 3 layers, molecular (ML), granular (GL), hilus (HL) the hippocampal region considered for cell count density experiments at postnatal day 3 and day 7. All images shown are 20 µm sections, scale bar 50 µm. The count included all three layers of DG as shown in the sample image.

We also examined the hilus and molecular layers of the DG sub-regions because of their important role in postnatal hippocampal formation as germinal zones. We found different densities of reactive astrocytes and microglia immunoreactivity in both the hilus and molecular layers in the control vs IVH and IVH vs SFN-DFN treated pups at postnatal day 3 (Figures 3B, C vs 3E, F and 3E, F vs 3H, I). Indeed, absolute astrocyte and microglial cell counts in the DG region (Figure 3N; includes the dentate gyrus 3 layers ^22^), were significantly higher in IVH compared with healthy controls (Figure 3J astrocyte, L microglia). Whereas, after SFN-DFN treatment with either low dose DFN (50mg/Kg) or high dose (100mg/Kg), both cell types were reduced significantly (Figure 3J astrocyte, & 3L microglia; p<0.05; n=5 in each group) at postnatal day 3. A similar comparison on postnatal day 7 remained significantly lower only at the high dose DFN (100mg/Kg; Figure 3K astrocytes, M microglia; p<0.05; n=5 in each group).

### 3.5 SFN-DFN treatment results in reduced neuronal cell death induced after IVH in the hippocampal/DG sub-region

RBC lysis toxicants may affect neurogenesis in the DG and cause glial changes due to active inflammation and oxidative stress. We found that immune reactivity for TUNEL positive cells and NeuN-TUNEL double positive cells were elevated in IVH as compared with no IVH controls (Figure 4A-C vs 4D-F), whereas after SFN-DFN treatment the immune signals for NeuN-TUNEL positive cell death at day 3 were reduced (Figure 4D-F vs 4G-I). We observed similar findings on day 7 as on day 3 in IVH as compared with no IVH controls (Supplementary Figure 2A-C vs 2D-F). SFN-DFN treatment suppressed immune signal for double positive NeuN-TUNEL immunoreactivity signal indicating less neuronal cell death at postnatal day 7 (Supplementary Figure 2D-F vs 2G-I).

**FIGURE 4:**
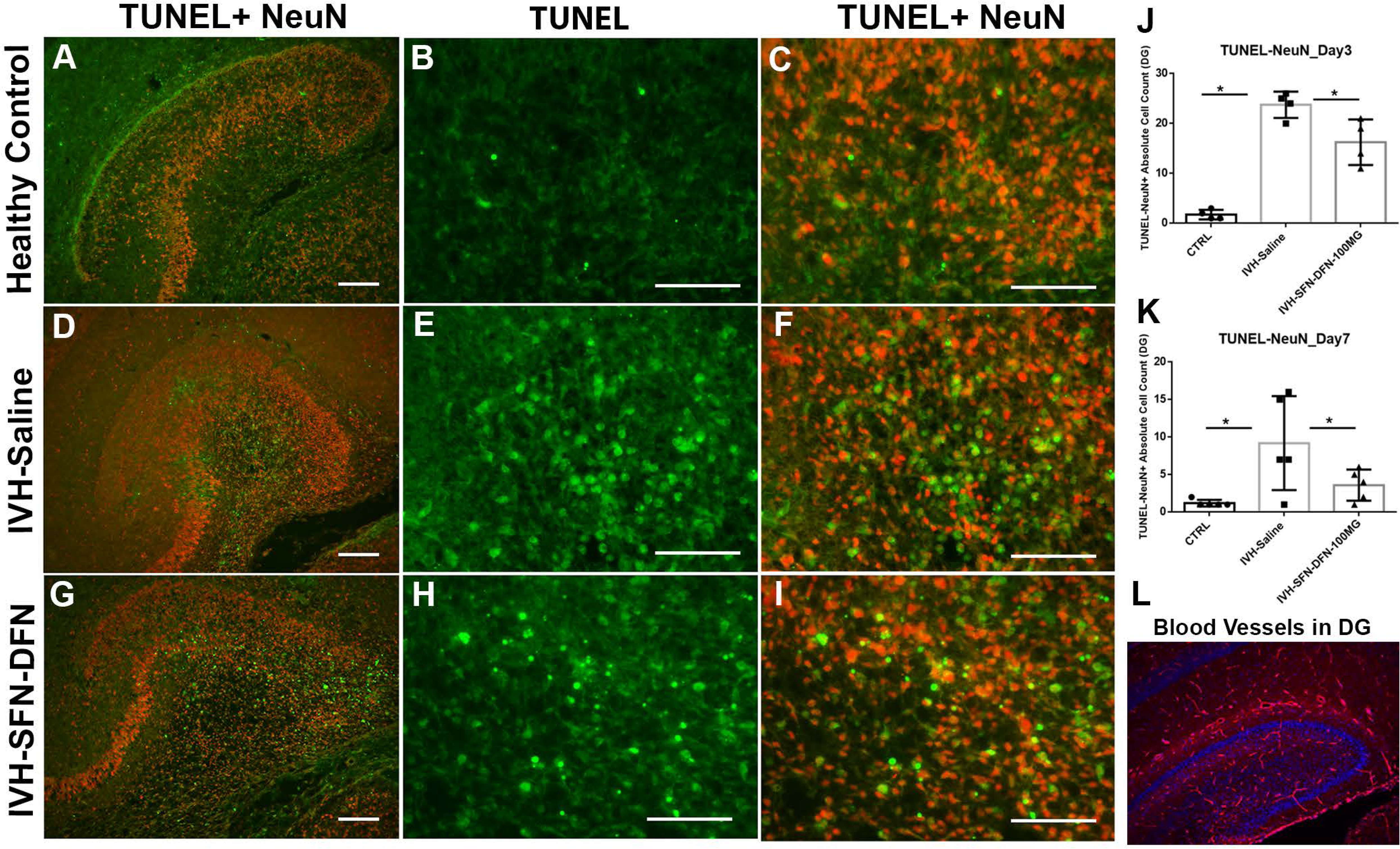
Sulforaphane and Deferoxamine treatment suppressed TUNEL positive neuronal cell death when co-localized with NeuN positive neurons during postnatal development of the hippocampal/DG region in premature rabbit pups at postnatal day 3. A-C) Representative immunofluorescence-stained images on coronal sections using TUNEL staining (positive for programmed cell death) combined with NeuN (neuronal marker) in the hippocampal/DG sub-regions in healthy control pups at postnatal day 3. Low magnification merged image 20X (A) of controls. High magnification 40X single stain for TUNEL (green), no immunoreactivity (B) and normal Neurons (NeuN, red) immunoreactivity in controls. D-F) Representative immunofluorescence-stained images on coronal sections using TUNEL staining (positive for cell death) combined with NeuN (neuronal marker) in the hippocampal/DG sub-regions in IVH premature pups at postnatal day 3. Low magnification merged image 20X (D). High magnification 40X single stain for TUNEL (green), shows positive immunoreactivity indicating overall programmed cell death (E) and overlapping with Neurons (NeuN, red) immunoreactivity indicating neuronal cell death (F). G-I) Representative immunofluorescence-stained images on coronal sections using TUNEL staining (positive for cell death) combined with NeuN (neuronal marker) in the hippocampal/DG sub-regions after SFN-DFN treated IVH premature pups at postnatal day 3. Low magnification merged image 20X (G). High magnification 40X single stain for TUNEL (green), shows fewer cells with positive immunoreactivity indicating reduced overall cell death (H) and no overlap with neurons (NeuN, red) immunoreactivity indicating reduced neuronal cell death (I). J-K) Scatter plot with bar graph shows cell density quantification for TUNEL + NeuN double positive cells after SFN-DFN treatment in coronal immunofluorescence-stained sections in premature rabbit pups at postnatal day 3 and day 7. The IVH pups showed significantly higher numbers of neuronal cell death; but, after SFN-DFN treatment this was significantly reduced in number. The data depicted is the cell count for 20X images showing the mean absolute cell count with S.D. Sample size of 5 in each group, *P < 0.05. L) Representative immunofluorescence-stained image of a coronal section using CD31 antibody (blood vessel marker) for the hippocampal region at postnatal day 3. The DG regions shows densely growing blood vessels. Image shows a 20 µm section.

Total NeuN-TUNEL double-positive cells in hippocampal coronal sections showed a significant increase cell density on day 3 and day 7 in IVH pups as compared to the no IVH control pups in the dentate gyrus (Figure 4J and Figure 4K, p<0.05 for both day 3, n=4 and day 7, n=5). We observed a significantly decreased NeuN-TUNEL positive cell density in the DG after SFN-DFN treatment compared to IVH with no SFN-DFN treatment at day 3 and day 7 (Figure 4J and Figure 4K, p<0.05 for both day 3, n=4 and day 7; n= 5 in each group).

### 3.6 Comparative transcriptomic analysis for multiple cell death types after hemorrhage and effects of SFN-DFN treatment

#### 3.6.1 Ferroptosis

Iron-dependent programmed cell death (i.e. ferroptosis) is associated with hemorrhagic CNS injuries where free iron activates apoptotic mechanism involving intracellular free iron. We set upon an *a priori* discovery approach to interrogate intracellular processes that we may have not previously hypothesized. and performed mRNA-seq assays to assess ferroptosis regulated gene expression profiles in our experimental groups at day 3. The gene count data were identified based on log2-fold change where a value of 1 indicates a 2-fold increase and -1 is a 50% reduction relative IVH vs control. The RNA-seq transcriptome analysis showed that a preponderance of gene transcripts had down regulated from control during early IVH injury at day 3 (Figure 5A &B) and then after SFN-DFN treatment fell further. The analytical list of up-and-down regulated ferroptosis-related genes are shown in the table (Figure 5C). For example, the genes XBP1 (endoplasmic reticulum stress response), GPX4 (glutathione peroxidase 4; uses glutathione to neutralize lipid hydroperoxides), NFE2L2 (i.e. NRF2; activates genes that produce antioxidants and detoxifying protective enzymes), FTH and FTL (cellular iron storage proteins and regulates toxicity), GCLM9 (a major antioxidant enzyme in the brain), SLC38A1 (glutamine uptake to detoxify ammonia and producing glutamate) and HMOX1 (a multifaceted function with heme breakdown and iron release) increased ∼64-fold (Fig 5A-C). Interestingly, those same genes: XBP1, GPX4, NFE2L2/NRF2, FTH, FTL, GCLM9, SLC38A1 and HMOX1 all showed a pattern of low expression in healthy controls but, following IVH increased and then returned toward lower levels after SFN-DFN treatment (5 A-C).

**FIGURE 5:**
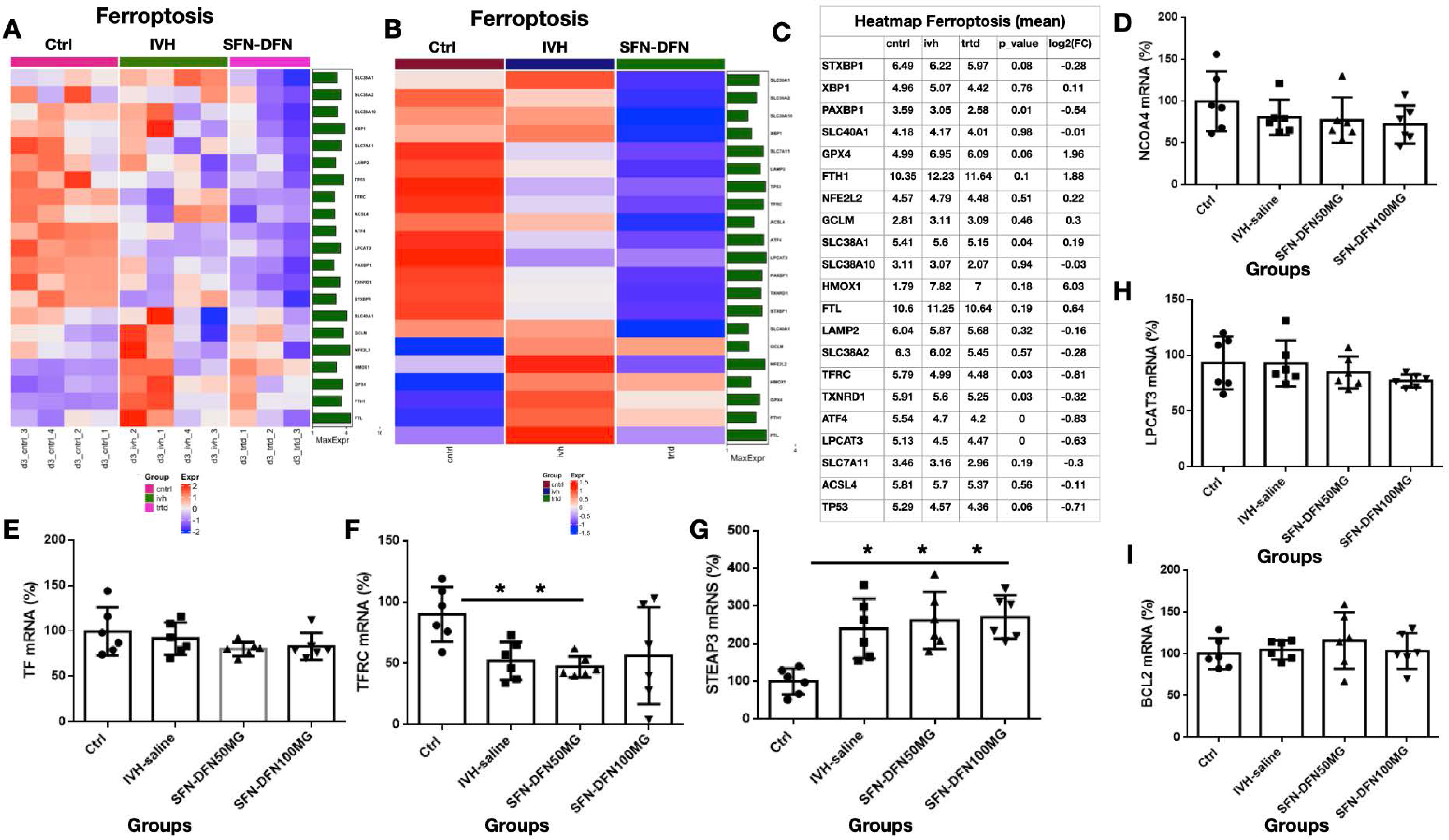
Heat maps of dysregulated genes belonging to functional proteins in hippocampal/DG tissue dissected RNA transcriptional profiling for ferroptosis in premature rabbit pup model of IVH and effects of SFN-DFN treatments at postnatal day 3. A-B) Representative heat map for ferroptosis differentially expressed genes (DEGs) and effect of SFN-DFN treatment at postnatal day 3. Sample wise heat map (A) and mean distribution DEG group wise (B). RNA isolated from dissected hippocampal/DG tissue. Data presented in Log2 mean fold change (n= 4 control, 4 IVH and 3 treated). C) Table showing key ferroptosis genes up-regulated and down-regulated (Log2 fold change) in three experimental groups. Scale bar indicates color coded changes (red to green). D-F) Scattered plot with bar graph showing TaqMan gene expression quantification using hippocampal/DG tissue dissected and isolated RNA in premature rabbit pups at day 3 and effects of SFN-DFN treatment. The mRNA expression for: E) Transferrin (TF), F) transferring receptor (TFRC), and G) STEAP3 showed TFRC was significantly decreased and STEAP3 was significantly increased after IVH. Sample size 6 in each group, P < 0.01 for CTRL vs IVH day3. D, H-I) Scatter plot with bar graph showing TaqMan gene expression quantification using hippocampal/DG tissue dissected and isolated RNA in premature rabbit pups at day 3 and effects of SFN-DFN treatment. The mRNA expression for: D) NCOA4, E) LPCAT3 and F) BCL2 showed no significant changes on day 3. Sample size 6 in each group.

Iron transport related protein regulated genes TF binds to Fe^3+^ and internalizes it into the cell via transferrin receptor-1 surface protein (TfR1) through an endocytosis process. TfR1 protein is a product of the TFRC gene transcript and showed a significant decrease but no changes in TF levels at early injury on day 3. This trend indicates highly localized intracellular response to iron accumulation rather than a systemic shift. The pattern of gene transcripts related to iron homeostasis, regulatory, and transport proteins (FTL, FTH and TFRC) indicated that there was an iron overload/clearance issue in the hippocampus after IVH.

The solute carrier family protein members (SLC) are involved in glutathione production (antioxidant). SLCs either: i) failed to change significantly (SLC1A5 (not shown); SLC39A14 (not shown), or ii) fell slightly, consistent with reducing intracellular export of iron after IVH and fell even further after SFN-DFN for SLC7A11 (brings cystine into cells in exchange for glutamate to increase glutathione antioxidant); SLC38A1 and SLC38A2 (both glutamine transporters); and SLC40A1 (transports iron into blood stream across cells), These SLC results indicated that iron exporting from cells was diminished and that antioxidant systems were impaired after IVH and even after SFN-DFN treatment.

We next assessed gene ontology (GO) enrichment analysis (categorizes gene expression into molecular, biological functional patterns, and cellular components). We assessed regulated genes (FTH, FTL, GCLM, GPX4 and HMOX1) in the following analysis. The GO biological process enrichment analysis data fell into intracellular iron homeostasis (GO:0006879), sequestering of metal ion (GO:0051238), responses to oxidative stress (GO:0006979), and mechanisms of iron ion transport (GO:0006826). Interestingly, the molecular functions of unregulated genes (not shown) that are also recognized to be involved in certain ferric and ferrous iron binding functions (GO:0008199; GO:0008198) and oxidoreductase (GO: 0016722), glutathione peroxidase (GO: 0004062) activities, taken together, indicated that RBC toxicants do not universally activate all related systems.

We used TaqMan gene expression assays to quantify the changes in key genes determined by RNAseq analysis. TaqMan assay results showed significantly reduced expression of TFRC (Figure 5F, n=6 in each group), which is a cellular iron uptake receptor (transferrin bound Fe^3+^ uptake) but a significantly increased level of STEAP3 (Metallo reductase that coverts Fe^3+^ into Fe^2+^) gene expression -which was unaffected by SFN-DFN (Figure 5G, p<0.05, n=6 in each group). This combination would allow Fe^3+^ accumulation extracellularly and Fe^2+^ formation inside the cell via increased STEAP3 activity. Of note the SFN-DFN treatment (100mg/dose) appeared to recover TFRC expression (Figure 6D-F, p<0.05, n=6 in each group). Expression of the other ferroptosis related network genes, TF, NCOA4 and LPCAT3 gene expression were not changed (Figure 5E, D, and H).

**FIGURE 6:**
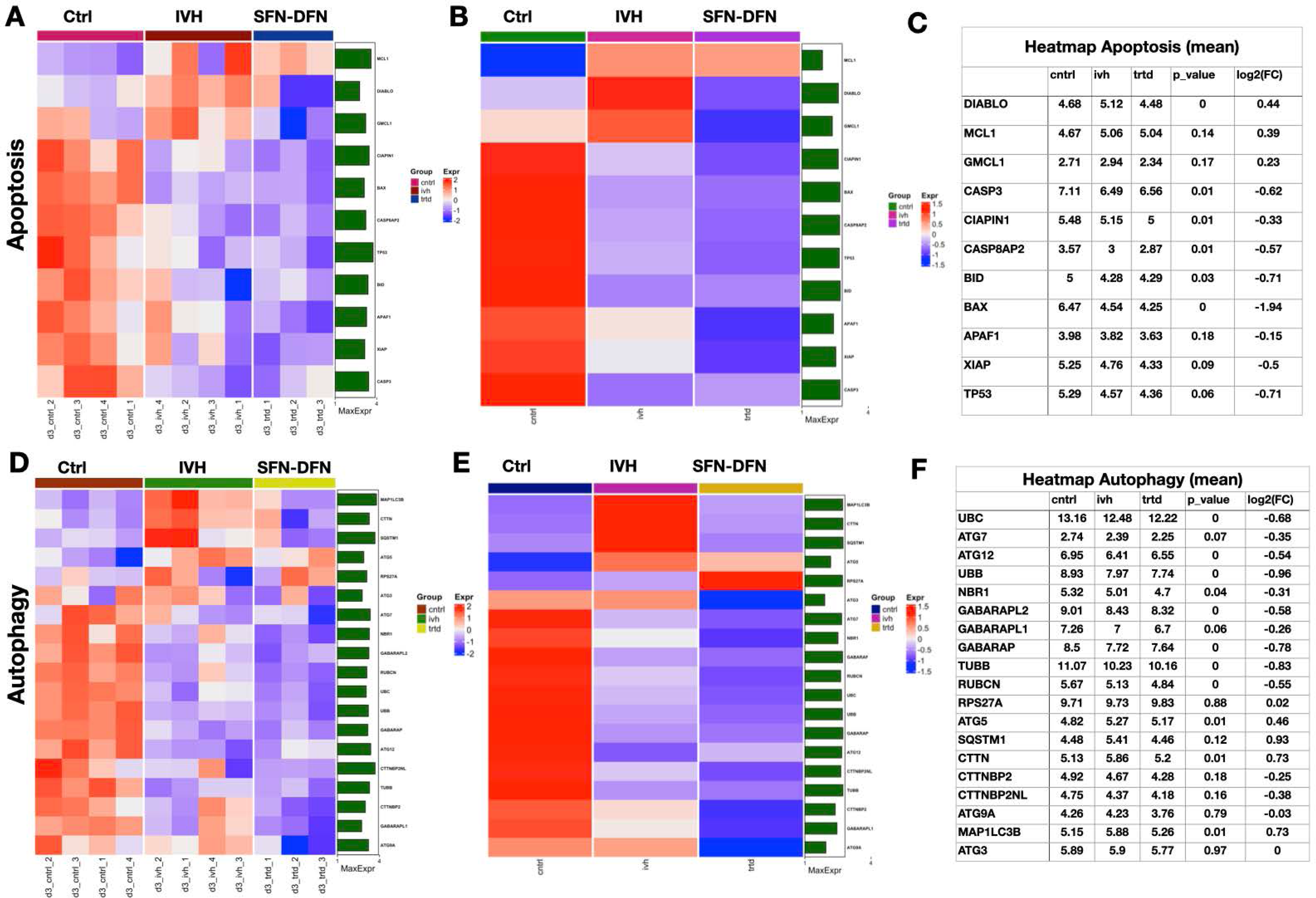
Heat maps of dysregulated genes belonging to functional proteins in hippocampal/DG tissue dissected RNA transcriptional profiling for apoptosis and autophagy genes in premature rabbit pup model of IVH and effect of SFN-DFN treatments at postnatal day 3. A-B) Representative heat map for apoptosis differentially expressed genes (DEGs) and effect of SFN-DFN treatment at postnatal day 3. Sample wise heat map (A) and mean distribution DEG group wise (B). RNA isolated from dissected hippocampal/DG tissue. Data presented in Log2 mean fold change. C) Table showing list of key apoptotic regulated genes up-regulated and down-regulated (Log2 fold change) in three experimental groups. D-F) Heat maps of dysregulated genes belonging to functional proteins in hippocampal/DG tissue dissected RNA transcriptional profiling for autophagy in premature rabbit pup model of IVH and SFN-DFN treatments at postnatal day 3. D-E) Representative heat map for autophagy differentially expressed genes (DEGs) and effect of SFN-DFN treatment at postnatal day 3. Sample wise heat map (A) and mean distribution DEG group wise (B). RNA isolated from dissected hippocampal/DG tissue. Data presented in Log2 mean fold change (n= 4 control, 4 IVH and 3 treated). F) Table showing list of key autophagy regulated genes up-regulated and down-regulated (Log2 fold change) in three experimental groups. Scale bar indicates color coded changes (red to green). Table showing key astrocyte regulated genes up-regulated and down-regulated (Log2 fold change) in three experimental groups.

#### 3.6.2 Apoptosis

On day 3, the DEG heat map indicated two clusters, The first cluster showed an increased gene expression after IVH that included DIABLO, MCL1 and GMCL1 (Fig 6 A-C). The MCL1 gene is an anti-apoptotic and member of BCL2 family of multiple function for apoptotic regulator group of proteins. And DIABLO is a mitochondrial secreted protein released into the cytosol that binds with an inhibitor of an apoptosis protein. A suppressed level of DIABLO, MCL and GMCL1 was observed after SFN-DFN treatment (Fig 6 A-C). The second cluster of genes with reduced expression included: CASP3, CIPIN1, CASP8AP2, BID, BAX, APAF1, XIAP and TP53. Caspase 3 and CASP8AP2 are mediators of programmed cell death during normal developmental pruning and neuronal degeneration. BAX promotes neuronal apoptosis and APAF1 is involved in forming the apoptosome complex that initiates the mitochondrial pathway of apoptosis. The X-linked inhibitor of apoptosis protein (XIAP) and TP53, cell protector genes were significantly decreased in IVH compared with healthy controls. After SFN-DFN treatment, there was no significant change in expression of any of these genes compared with IVH without treatment at day 3 (Fig 6A-C). We confirmed this by assessing a key anti-apoptotic protein BCL2 using real-time TaqMan assay and found no difference between experimental groups (Fig 5 I, n=6 samples in each group).

#### 3.6.3 Autophagy

Autophagy related genes (ATG) maintain neuronal health by removing damaged and misfolded proteins to further regulate and suppress inflammation. We observed that ATG related gene transcripts ATG3, AGT7, AGT9A, AGT12 and NBR1 were at reduced levels in their expression after IVH but not changed after SFN-DFN treatment (Figure 6D-F). In contrast, gene expression of ATG5, SQSTM1 and CTTN were all increased. These results indicate that autophagy type of cell death is relatively, less engaged after IVH and although there was a further trending downward after SFN-DFN (Figure 6F) the effects were minimal at day 3.

To summarize RNAseq data on cell death process mainly involved ferroptosis mechanisms and to some extent apoptotic cell death due to multiple dysregulated gene transcripts in the hippocampal/DG region following IVH. Further, SFN-DFN treatment limited activation of caspases and programmed cell death in the hippocampal/DG region. Most of the autophagy genes were neither activated in this region after IVH nor affected by administration of SFN-DFN.

### 3.7 Comparative transcriptome analysis for DEG associated with astrocyte and microglia infiltration

Astrocytes and microglia in the CNS are the first to respond to hemorrhagic toxicants within hours and induce the release of pro-inflammatory molecules that can exacerbate hippocampal damage (Figure 7 A-C). Up-regulated genes in astrocytes after IVH were: GFAP, ALDHL1, GPC5, GLUL, GLDC, CCL2/MCP1 and CD44. The down regulated genes were SOX9, AQP4, KCJN10, GJA1, GJA10 NFIA, FABP7 and DiO2 (Figure 7C, table). These genes are all involved in gliogenesis (ID GO: 0042063) and glial differentiation (ID GO: 0010001). Interestingly the reduced expression of NFIA, SOX9 is consistent with dysregulated glial cell fate in the injured hippocampus (ID GO:0021781). The molecular functions of genes GJA1 and GJA10 are involved in gap junction pore channel activity (ID GO: 0005243 and 0022829). The genes AQP4, GFAP, GLUL, KCNJ10 (ID GO: 0097386) and GJA1 were involved in glial cell projection and connexin complex formation (ID GO:0005921).

**FIGURE 7:**
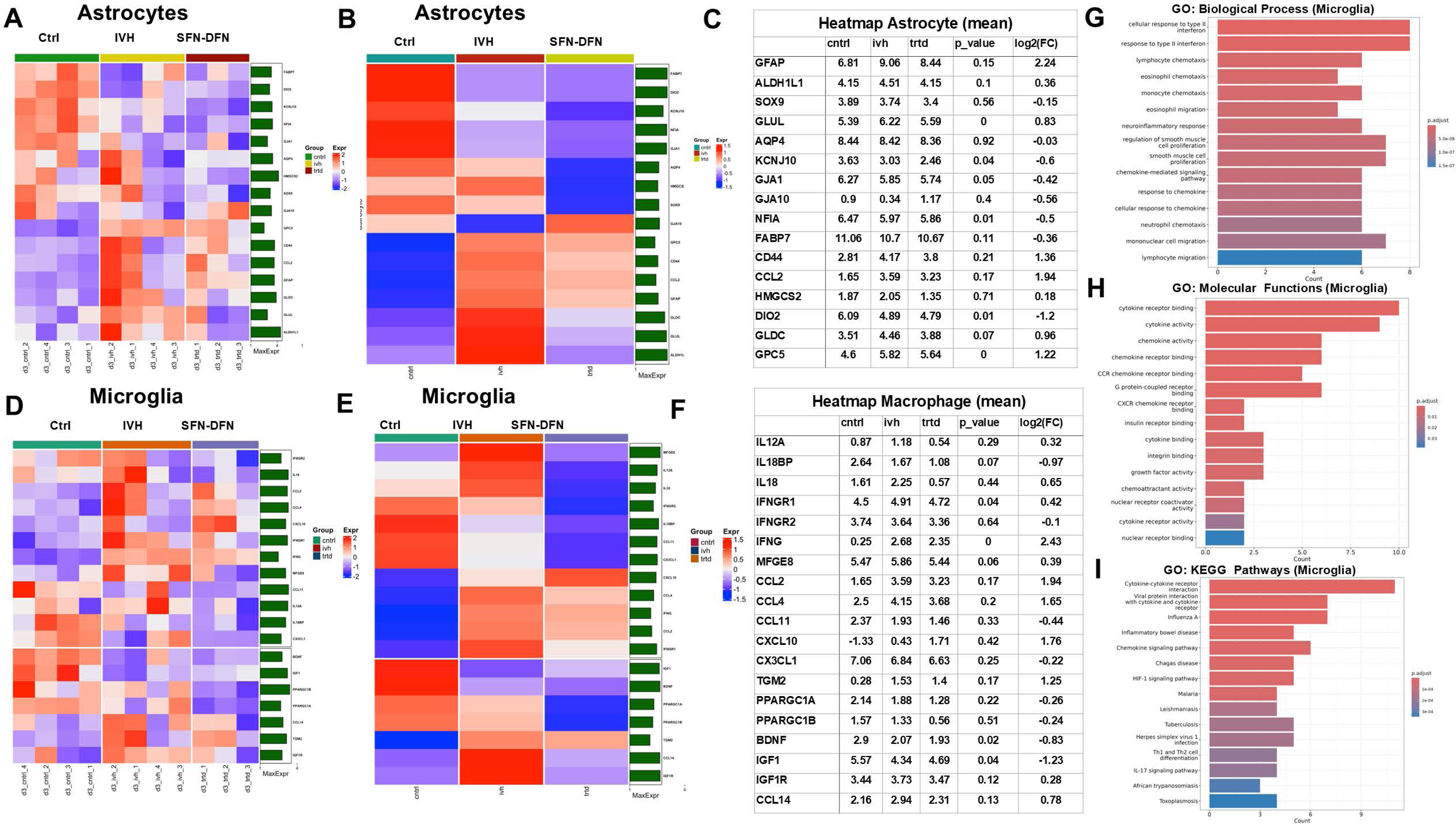
Representative heat map for astrocyte-and microglia-related differentially expressed genes (DEGs) and effect of SFN-DFN treatment at postnatal day 3. A-B) Representative heat map for astrocyte differentially expressed genes (DEGs) and effect of SFN-DFN treatment at postnatal day 3. Sample wise heat map (A) and mean distribution DEG group wise (B). RNA isolated from dissected hippocampal/DG tissue. Data presented in Log2 mean fold change (n= 4 control, 4 IVH and 3 treated). Scale bar indicates color coded changes (red to blue). C) Table showing list of key astrocyte-related genes up-regulated and down-regulated (Log2 fold change) in three experimental groups. D-E) Representative heat map for microglia-related differentially expressed genes (DEGs) and effect of SFN-DFN treatment at postnatal day 3. Sample wise heat map (D) and mean distribution DEG group wise (E). RNA isolated from dissected hippocampal/DG tissue. Data presented in Log2 mean fold change (n= 4 control, 4 IVH and 3 treated). Scale bar indicates color coded changes (red to blue) F) Table showing list of key microglia regulated genes up-regulated and down-regulated (Log2 fold change) in three experimental groups. G-I) Gene set enrichment analysis of microglia un/down regulated sets of genes involved in specific biological (G), molecular, and KEGG pathway functions (I). Key microglia regulated genes up-regulated and down-regulated (Log2 fold change) in three experimental groups.

In microglia (Figure 7 D-F), up-and-down regulated genes are shown in the table (Figure 7F, table). The pro-inflammatory cytokine IL12A and IL18 were over expressed after IVH but reduced after SFN-DFN treatment (Figure 7F, table). After 12-24 hours post-injury, activated microglia are the principle source of IL18 production.^23^ Significantly, these increases were reduced after SFN-DFN treatment.

IFNγ (interferon gamma) is a key inflammatory molecule and IFNγR1, it’s receptor, contribute to microglia polarization into the pro-inflammatory M1 phenotype; ^24^ this shift results in reduced phagocytotic function delaying hemorrhagic debris removal. ^25, 26^ Importantly, IFNγ also stimulates chemokines that serve a key role in recruiting T cells, NK cells and monocytes to sites of inflammation including: CXCL9, CXCL10 and CCL2 (MCP-1) -production particularly from astrocytes and microglia. ^24^ Chemokines and chemokine receptors CCL2, CCL4 and CCL14 are pro-inflammatory after hemorrhage ^27^ and each was over expressed in IVH compared with controls; whereas, SFN-DFN treatment brought them down. On other hand the transcripts of genes: BDNF, PPARGC1A, PPARGC1B, CCL11and CX3CL1 were reduced after IVH and decreased even further after treatment with SFN-DFN. In contrast, CXCL10 fell after IVH but rose to supranormal levels after SFN-DFN. IGF1 was reduced in expression after IVH but increased after SFN-DFN (Fig 7F).

We next assessed gene ontology (GO) enrichment analysis (categorizing gene families by molecular, biological functions and cellular components). Microglia associated genes (Figure 7G, table), were primarily connected to cellular responses to type-II interferon, lymphocyte and monocyte chemotaxis (8/19 genes for both, IDs, G: 0071346 and GO: 00034341. Major genes in the microglia network were connected to cytokine activity and cytokine receptor binding functions (IDs, GO :0005126; 10/19 and GO: 0005125; 9/19); other genes are reported in the data base (Figure 7H). An expanded list of genes with molecular functions is presented in Supplemental Table 1 (Appendix). We next explored KEGG pathway (Kyoto Encyclopedia of Genes and Genomes database) enrichment analysis to identify which genes are changed and fall into known KEGG pathways. Our data directed us to cytokine-cytokine receptor interaction, HIF-1 signaling pathways and multiple other immune signaling pathways (Figure 7I), all consistent with accelerated inflammatory mechanisms.

## 4. DISCUSSION

Intraventricular hemorrhage is a common problem of premature infants and is associated with white matter injury, neurodevelopment and cognitive disabilities persisting for years after premature birth. ^28^ Despite substantial research efforts, currently, no effective preventive or disease modifying clinical treatment for neonatal IVH can recover the brain to normal development. One prior notable and encouraging exception is the DRIFT protocol (Drainage, Irrigation, and Fibrinolytic Therapy). The study demonstrated modest long-term benefits in human neonates but was halted prematurely due to the risk of secondary perinatal hemorrhage. ^29, 30^ The findings of the DRIFT study underscored the importance of a multitiered approach directed at interventions that concurrently address early hematoma/iron clearance to attenuate later injurious inflammatory-mediated injury arising from hemolysis-derived toxic mediators.

In this report we sought to test a minimally invasive pharmacological alternative to expand upon that tiered approach concept by using a relevant animal model of endogenous bleeding. Most rodent models using injected exogenous whole blood and blood components were informative regarding the impact of RBC debris causing neuroinflammation and reduced neurogenesis, but they fail to mimic the more complex human condition where *spontaneous* hemorrhage into the developing forebrain arises from rupture of immature vasculature. The spontaneous bleeding also creates a mass effect further reducing nutrient and oxygen delivery. Moreover, these events are likely to affect other periventricular regions like the hippocampal dentate germinal region, yet, the hippocampus has received limited attention as a target structure. Therefore, to address this gap, we used a premature rabbit model of IVH in which intraperitoneal glycerol-induces an osmotic injury that causes a reproducible, spontaneous post-hemorrhagic insult in 70% of the pups. ^31, 32^

Using the rabbit pup model, we recently reported that periventricular accumulation of free hemoglobin and iron occurs after IVH and that pharmacologic agents can suppress this accumulation and improve recovery by attenuating the cascade of inflammatory responses arising from extravasation and lysis of red blood cells.^19^ In the present study, using the same premature rabbit model of IVH, we found that: i) hemorrhagic injury debris occurs throughout the entire hippocampus (Figure-1), ii) evidence for ferroptosis as primary mode of cell death rather than predominance of caspase-mediated apoptosis or autophagy (at early injury day 3; Figures 2, 4–6), and most importantly, iii) SFN-DFN reduced inflammatory markers as evidenced by reduced reactive astrocytes with lower expression of GFAP mRNA (Figures 2P and 7C, table); as well as fewer activated microglia. SFN-DFN (100mg dose). In addition, we also found lower steady-state levels of hemeoxygenase 1 mRNA (HMOX1; a Nrf2 regulated gene whose product is involved in breaking down free heme protein); this result was associated with a reduction in free hemoglobin levels (Figures 2Q, 2M and 5C, table), ^33^ and lower inflammatory molecules. Taken together, our results offer a *proof-of-concept* that SFN-DFN can abrogate multiple offending pathways including altered gene expression involving ferroptotic cell death and inflammation after IVH. This suggests that the combined effects of SFN-DFN treatment will likely contribute to a microenvironment conducive to recovery of normal brain development.

### Glial Cell Activation and neuroinflammation

Morphological changes of astrocytes and microglia are well characterized as evidence of CNS injury and inflammation. Previous studies identified free hemoglobin as a potent inducer of pro-inflammatory markers in the SVZ and hippocampus.^34, 35^ In the present hippocampal study, we also identified a morphological transition toward activated astrocytes and microglia in the molecular and hilus region of the DG (Figure 3); an area with significant hemoglobin and free iron deposition (Figure 2).^12^ Of note, in this study, we observed strong GFAP immune reactivity and mRNA expression in IVH, whereas SFN-DFN treatment reduced astrocyte immune reactivity in the hilus (Figure 3). Surprisingly, we also identified a regional specificity to these effects as no astrogliosis was observed in hippocampal CA1 to CA4 regions after IVH or when compared with healthy controls (Figure 2). In the astrocyte transcriptomic data, SFN-DFN reduced over expression of GFAP and CCL2/MCP1 and down regulated NFIA, SOX9 and KCNJ10 transcripts which are all involved in gliogenesis (ID GO: 0042063) and glial differentiation (ID GO: 0010001). In the microglial transcriptome, the pro-inflammatory cytokines IL12A and IL18 were over-expressed in the injured hippocampus ^23^ and were reduced after combined treatment as well (Figure 7 F).

Although a Nrf2 regulated gene, SFN-DFN treatment had little effect on the elevation of heme oxygenase RNA (Figure 2Q; which detoxifies free heme into biliverdin and then converts that to bilirubin for excretion along with carbon monoxide and Fe^++^, which is recycled for hematopoiesis); this may indicate it is already maximally expressed. HEMOX-1 is among several other Nrf2 responsive genes like GPX, GCLM (both increased in IVH and stayed at higher levels even after treatment), SLC38A1, SLC40A1 (no significant changes), ACL4 and TP53 (slightly decreased in IVH and similarly even with treatment) (Figure 5), and are considered protective against oxidation and inflammation -consistent with the interpretation that the scope of responses to SFN-DFN are complementary to ongoing endogenous processes.

### Hippocampal RBC Lysis Products and Cell Death

On postnatal day 3 (early phase injury), on histology, iron was deposited in the hippocampal/DG parenchyma and was elevated in the CSF (Figure 2); SFN+DFN decreased iron in both hippocampal tissue and in circulating CSF (Figure 2E-H vs 2I-L for tissue and Figure 2N for CSF); however, although reduced slightly, tissue free iron concentrations did not achieve statistical significance in hippocampal lysates (Figure 2-O). Elevated free hemoglobin was reduced by SFN-DFN treatment (Figure 2M).

Cell death was prominent (TUNEL-NeuN double positive cells) in brain regions associated with large quantities of iron deposition (Figure 2), such as in the dentate hilar area and the molecular layer (Figure 2 & 4). Moreover, transcriptome data from hippocampus/DG-dissected tissue RNA showed elevated gene transcripts after IVH for iron storage proteins FTL (light ferritin subunit; storage of iron) and FTH (heavy ferritin subunit which participate in converting highly reactive Fe^2+^ into safer Fe^3+^ via ferroxidase activity. ^36, 37^ These mRNAs were reduced by SFN-DFN, consistent with improved iron clearance. In contrast, iron binding transferrin (TF) was unchanged while TFRC (transferrin receptor protein; cellular iron uptake) was reduced which is indicative of excessively loaded intracellular iron regulatory storage proteins (FTL and FTH).^38^ In contrast, gene transcripts GPX4 (glutathione peroxidase 4; reduces ferroptosis by preventing harmful lipid peroxide accumulation by conversion to lipid alcohols), HMOX1 and NRF2 remained elevated during IVH even after SFN-DFN suggestive of an already maximally activated protection from iron-induced free radical toxicity and cell death.

#### Ferroptotic Cell Death

Transferrin (TF) mRNA was unchanged after IVH, while transferrin receptor (TFRC) mRNA expression was suppressed after IVH consistent with known exosome mediate phagocytosis mechanisms along with elevation of STEAP3 mRNA, a transcript for an intracellular free iron recycling protein involved in programed cell death known as *ferroptosis* (Figure 4–5). ^37, 39, 40^. We interpret this to mean that a decrease in TFRC cell membrane protein will reduce cell uptake of extracellular transferrin (TF) bound iron leaving TF levels unchanged. However, after IVH an increase in STEAP3 function will reduce ferric iron (Fe^3+^) to a more reactive ferrous form (Fe^2+^) fueling the Fenton Reaction creating hydroxyl free radicals (OH.). ^41, 42^ This, in turn, leads to accumulation of toxic lipid ROS and cell injury

In the CSF, SFN-DFN treatments were associated with a large reduction in extracellular iron and free hemoglobin on day 3 (early injury) (Figure 2 M, N vs O); yet, there was no independent effect of SFN-DFN treatment on iron clearance in tissue lysates or on STEAP3 mRNA levels. TFRC increased to baseline after the 100mg (high dose DFN) consistent with enhanced removal of iron. Presumably, the reduced CSF iron should progressively attenuate its impact on intracellular dysfunction and ferroptosis as supported by evidence of reduced TUNEL-positive cells we identified.

#### Apoptotic cell death

IVH is associated with slightly elevated expression of the genes DIABLO, MCL1 and GMCL1 (germ cell-less-1); SFN-DFN treatment suppressed this response except for MCL1 (myeloid leukemia-1; Figure 6A-C). This is significant since the MCL1 gene is anti-apoptotic and a member of BCL2 family of multiple function, apoptotic regulator group proteins. It indicates that this endogenous homeostatic response is not impaired by SFN-DFN treatment and may in fact, help amplify MCL1’s impact on reducing apoptosis. Moreover, after IVH, neither the mRNA for key enzymes involved in caspase**-**mediated apoptotic cell death (NCOA4, LPCAT3, BCL2) were affected after IVH nor did treatment with SFN-DFN alter their *status quo* (Figure 6A-C).

#### Autophagy

Another important cellular homeostatic system where, for several key genes involved (Figure 6 D-F), the transcriptome data showed less than a 2-fold change from baseline mRNA thus, not meeting a conventional definition of significant alteration in gene expression. Taken together, we interpret the patterns of change in these three key mechanisms (sp. ferroptosis, apoptosis, autophagy) as compelling in implicating ferroptosis as the principal mechanism of cell death after IVH in the hippocampus. This supports our contention that an iron chelator is likely to attenuate the progression of post-injury inflammatory processes becoming an important strategic insight into the pathophysiologic response to IVH in the hippocampus.

Overall, SFN-DFN treatment shows promise in reducing inflammation in the hippocampus. Recovery in the DG germinal zone is particularly important as this region gives rise to new neuronal precursors that integrate into the hippocampal/DG formation during postnatal development and throughout life. The limitations of this study include the need for further confirmation of our mRNA transcriptome observations by quantifying changes in protein levels or their functions. In several cases, we already validated key genes by TaqMan PCR to add to the interpretation of the results (Figure 4 D-I). Changes in transcriptome expression for the many other genes were not categorically addressed as they were beyond the scope of this report; the key total gene list in the study is available in Supplementary Table 2 and raw transcriptome data available GEO-NCBI (GSE 305615). Future mechanistic studies on iron transport via the TF-TFRC-STEAP system as a means to generate a labile pool of less toxic ferrous and ferric iron forms may aid in reducing oxidative free radical damage. ^38^

#### In conclusion

SFN-DFN treatment reduced key pathological features of IVH including iron accumulation and ferroptosis, as well as fewer reactive astrocytes and reduced microglial infiltration. This was associated with a gene profile favoring ferroptotic cell death rather than apoptosis or autophagy pathways. Taken together, SFN-DFN treatment aided in creating a less hostile microenvironment. Moreover, the early phase decrease in accumulation of toxic byproducts such as free hemoglobin and iron that mitigated CNS injury may also enhance the survival of future therapies using transplanted stem cells that can adapt to the changing tissue microenvironment, reducing inflammation as “*living cellular therapy*.” ^43^

## Acknowledgements

The authors thank Dr. Carl Thompson, PhD, for helping statistical analysis. We acknowledge Rita Daly (Newborn Medicine, Boston children Physicians Group) for secretarial support and Jeanette Aprile in formatting and editing the manuscript for the Journal submission.

## Data Availability

The data generated from all experiments, which support the findings presented in this study, are available from the corresponding authors upon request, in accordance with institutional policies.

## AUTHOR CONTRIBUTIONS

GV, EFL: Conception, design, experiments, assembly, analysis, interpretation of the data and wrote the manuscript.

MS, CL, VCD, FH: Rabbit tissue and other sample collection, processing and performed experiments. MS, wrote first draft of the manuscript.

JV, LV, SSH, SS, RS: Performed RNA-Seq and data analysis and manuscript review and editing, comments and suggestions.

## SUPPLEMENTARY MATERIAL

**SUPPLEMENTARY FIGURE 1:** Postnatal hippocampal/DG region and hematoma/debris accumulation after intraventricular hemorrhage in premature rabbit pups.

A-D) Representative hematoxylin and eosin staining (H&E) on coronal serial section images showing hippocampal/DG sub-regions in healthy control premature rabbit pups without hemorrhage at postnatal day 7.

E-F) Representative H&E staining on coronal serial section images showing large hematoma accumulation in the hippocampal/DG regions in premature rabbit pups with morphologically damaged DG layers at day 7.

Every 10th sections was taken from anterior to posterior region for H&E stained sections and stitched images were taken at low magnification using KEYENCE microscope (Keyence corporation of America, Illinois, USA). The scale bar for all images 500 µm for stitched images.

**SUPPLEMENTARY FIGURE 2**: Sulforaphane and Deferoxamine treatment suppressed TUNEL positive cell death and its co-localization with NeuN positive neurons during postnatal development of hippocampal/DG region in premature rabbit pups at postnatal day 7.

A-C) Representative immunofluorescence-stained images on coronal sections using TUNEL staining (positive for cell death) combined with NeuN (neuronal marker) in the hippocampal/DG sub-regions in healthy control pups at postnatal day 7. Low magnification merged image 20X (A). High magnification 40X single stain for TUNEL (green), no immunoreactivity (B) and normal Neurons (NeuN, red) immunoreactivity.

D-F) Representative immunofluorescence-stained images on coronal sections using TUNEL staining (positive for cell death) combined with NeuN (neuronal marker) in the hippocampal/DG sub-regions in IVH premature pups at postnatal day 7. Low magnification 20X merged image (D). High magnification 40X single stain for TUNEL (green), positive immunoreactivity indicated active cell death (E) and overlapping with Neurons (NeuN, red) immunoreactivity indicating neuron cell death (F).

G-I) Representative immunofluorescence-stained images on coronal sections using TUNEL staining (positive for cell death) combined with NeuN (neuronal marker) in the hippocampal/DG sub-regions after SFN-DFN treated IVH premature pups at postnatal day 3. Low magnification 20X merged image (G). High magnification 40X single stain for TUNEL (green), shows fewer immunoreactive positive cells indicating reduced cell death (H) and no overlap with neuronal (NeuN, red) immunoreactivity indicating fewer neuronal cell deaths (I).

**SUPPLEMENTARY TABLE 1:**

Table showing list of total number dysregulated genes in microglial and their connections for molecular functions.

**SUPPLEMENTARY TABLE 2:**

Expanded gene names for list of gene abbreviations used in the study.

