## Supplement Figures and Tables for "Combined pharmacotherapy suppress ferroptosis and inflammation after neonatal intraventricular hemorrhage"

**Supplementary Figure 1**

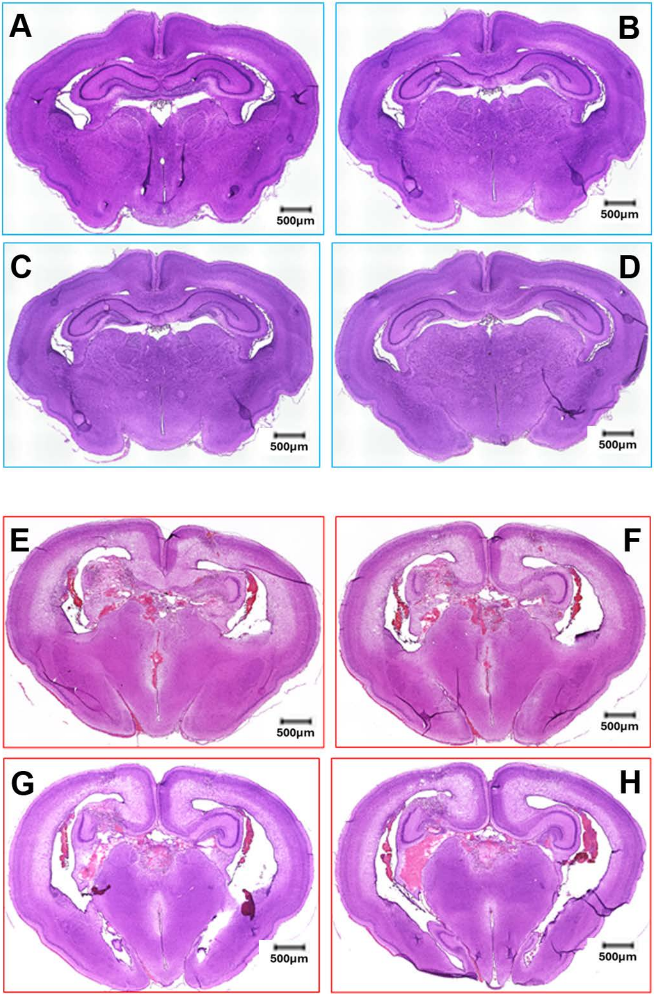

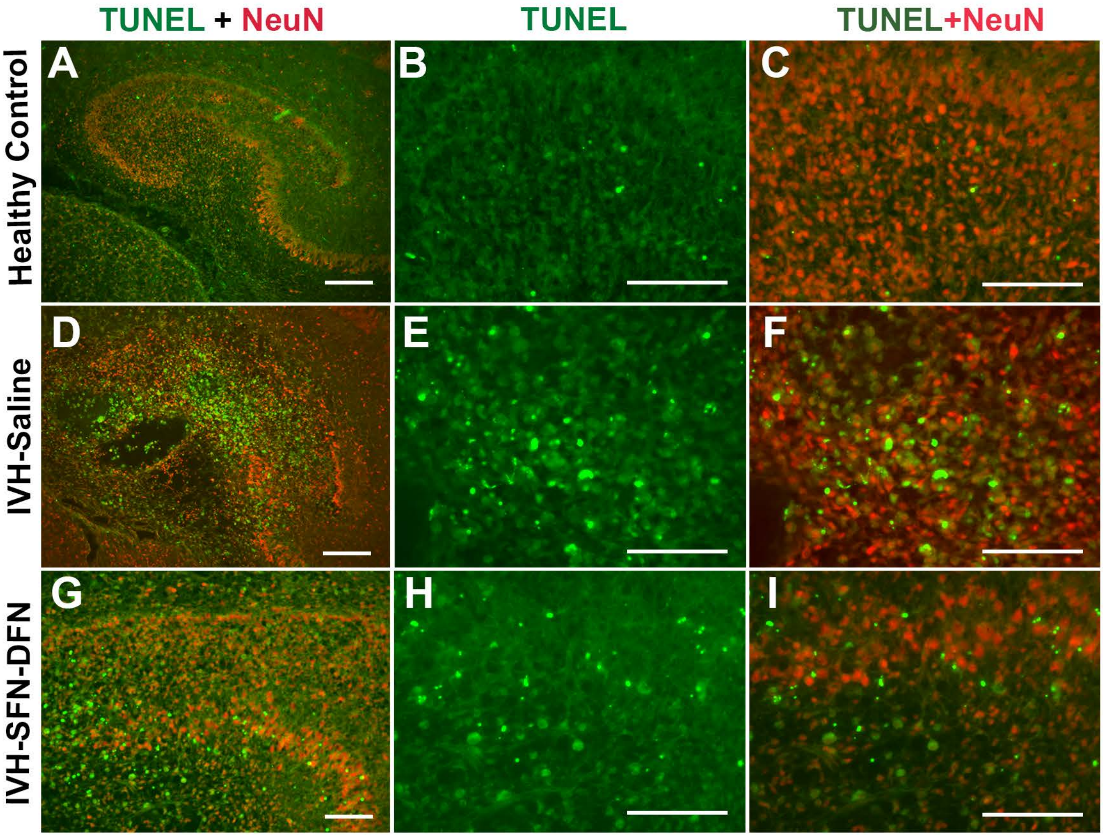

TUNEL + NeuN
TUNEL
TUNEL+NeuN
Healthy Control
IVH-Saline
IVH-SFN-DFN
A
B
C
D
E
F
G
H
I

**GO Molecular Function**

| ID | Description | GeneRatio | BgRatio | pvalue | p.adjust | qvalue | Gene ID | Count |
|---|---|---|---|---|---|---|---|---|
| **GO:0005126** | cytokine receptor binding | 10/19 | 273/18496 | 3.42114227577589E-14 | 2.29216532476984E-12 | 1.15238476657714E-12 | BDNF/CCL11/CCL14/CCL2/CCL4/CX3CL1/CXCL10/IFNG/IL12A/IL18 | 10 |
| **GO:0005125** | cytokine activity | 9/19 | 237/18496 | 6.6102250922672E-13 | 2.21442540590951E-11 | 1.11330106817132E-11 | CCL11/CCL14/CCL2/CCL4/CX3CL1/CXCL10/IFNG/IL12A/IL18 | 9 |
| **GO:0008009** | chemokine activity | 6/19 | 49/18496 | 6.65364448826736E-12 | 1.48598060237971E-10 | 7.47075872366861E-11 | CCL11/CCL14/CCL2/CCL4/CX3CL1/CXCL10 | 6 |
| **GO:0042379** | chemokine receptor binding | 6/19 | 74/18496 | 8.68233136050217E-11 | 1.45429050288411E-09 | 7.31143693515972E-10 | CCL11/CCL14/CCL2/CCL4/CX3CL1/CXCL10 | 6 |
| **GO:0048020** | CCR chemokine receptor binding | 5/19 | 50/18496 | 1.32822495173453E-09 | 1.77982143532427E-08 | 8.94804178010628E-09 | CCL11/CCL14/CCL2/CCL4/CX3CL1 | 5 |
| **GO:0001664** | G protein-coupled receptor binding | 6/19 | 291/18496 | 3.29013875714769E-07 | 3.67398827881492E-06 | 1.84709544260923E-06 | CCL11/CCL14/CCL2/CCL4/CX3CL1/CXCL10 | 6 |
| **GO:0045236** | CXCR chemokine receptor binding | 2/19 | 18/18496 | 0.00015147012597082 | 0.00144978549143499 | 0.000728878801664096 | CX3CL1/CXCL10 | 2 |
| **GO:0005158** | insulin receptor binding | 2/19 | 22/18496 | 0.000228130484714972 | 0.00191059280948789 | 0.000960549409326198 | IGF1/IGF1R | 2 |
| **GO:0019955** | cytokine binding | 3/19 | 145/18496 | 0.000417060240222063 | 0.0031047817883198 | 0.0015609272148662 | IFNGR1/IL12A/IL18BP | 3 |
| **GO:0005178** | integrin binding | 3/19 | 154/18496 | 0.000497337498763047 | 0.00333216124171241 | 0.00167524210109658 | CX3CL1/IGF1/MFGE8 | 3 |
| **GO:0008083** | growth factor activity | 3/19 | 162/18496 | 0.000576502837570812 | 0.00351142637429495 | 0.0017653675408867 | BDNF/IGF1/IL12A | 3 |
| **GO:0042056** | chemoattractant activity | 2/19 | 41/18496 | 0.000800446512235864 | 0.00446915969331691 | 0.00224686740276734 | CX3CL1/CXCL10 | 2 |
| **GO:0030374** | nuclear receptor coactivator activity | 2/19 | 55/18496 | 0.00143722251298703 | 0.00740722372077932 | 0.00372397736158583 | PPARGC1A/PPARGC1B | 2 |
| **GO:0004896** | cytokine receptor activity | 2/19 | 96/18496 | 0.00430404381486696 | 0.020597923971149 | 0.0103555941410333 | IFNGR1/IFNGR2 | 2 |
| **GO:0016922** | nuclear receptor binding | 2/19 | 137/18496 | 0.00857576039865447 | 0.03830506311399 | 0.0192578479127679 | PPARGC1A/PPARGC1B | 2 |
| **GO:0140375** | immune receptor activity | 2/19 | 147/18496 | 0.0098182725434521 | 0.0411140162757056 | 0.0206700474598991 | IFNGR1/IFNGR2 | 2 |
| **GO:0005165** | neurotrophin receptor binding | 1/19 | 12/18496 | 0.0122612075186457 | 0.0444763422087473 | 0.0223604548417897 | BDNF | 1 |
| **GO:0008603** | cAMP-dependent protein kinase regulator activity | 1/19 | 12/18496 | 0.0122612075186457 | 0.0444763422087473 | 0.0223604548417897 | CXCL10 | 1 |
| **GO:0031994** | insulin-like growth factor I binding | 1/19 | 13/18496 | 0.0132765200623126 | 0.0444763422087473 | 0.0223604548417897 | IGF1R | 1 |
| **GO:0043560** | insulin receptor substrate binding | 1/19 | 13/18496 | 0.0132765200623126 | 0.0444763422087473 | 0.0223604548417897 | IGF1R | 1 |

**Supplementary Table 2**. Expanded gene names for list of gene abbreviations used in the study.

| | |
|---|---|
| STXBP1 | Syntaxin binding protein 1 |
| XBP1 | X-box binding protein 1 |
| PAXBP1 | PAX3 and PAX7 binding protein 1 |
| SLC40A1 | Solute carrier family 40 member 1 |
| GPX4 | Glutathione peroxidase 4 |
| FTH1 | Ferritin heavy chain 1 |
| NFE2L2 | NFE2 like bZIP transcription factor 2 |
| GCLM | Glutamate-cysteine ligase modifier subunit |
| SLC38A1 | Solute carrier family 38 member 1 |
| SLC38A10 | Solute carrier family 38 member 10 |
| HMOX1 | Heme oxygenase 1 |
| FTL | Ferritin light chain |
| LAMP2 | Lysosomal associated membrane protein 2 |
| SLC38A2 | Solute carrier family 38 member 2 |
| TFRC | Transferrin receptor |
| TXNRD1 | Thioredoxin reductase 1 |
| ATF4 | Activating transcription factor 4 |
| LPCAT3 | Lysophosphatidylcholine acyltransferase 3 |
| SLC7A11 | Solute carrier family 7 member 11 |
| ACSL4 | Acyl-CoA synthetase long chain family member 4 |
| TP53 | Tumor protein p53 |
| DIABLO | Diablo IAP-binding mitochondrial protein |
| MCL1 | MCL1 apoptosis regulator BCL2 family member |
| GMCL1 | Germ cell-less 1 spermatogenesis associated |
| CASP3 | Caspase 3 |
| CIAPIN1 | Cytokine induced apoptosis inhibitor 1 |
| CASP8AP2 | Caspase 8 associated protein 2 |
| BID | BH3 interacting domain death agonist |
| BAX | BCL2 associated X apoptosis regulator |
| APAF1 | Apoptotic peptidase activating factor 1 |
| XIAP | X-linked inhibitor of apoptosis |
| TP53 | Tumor protein p53 |
| UBC | Ubiquitin C |
| ATG7 | Autophagy related 7 |
| ATG12 | Autophagy related 12 |
| UBB | Ubiquitin B |
| NBR1 | NBR1 autophagy cargo receptor |

| | |
|---|---|
| GABARAPL2 | GABA type A receptor associated protein like 2 |
| GABARAPL1 | GABA type A receptor associated protein like 1 |
| GABARAP | GABA type A receptor-associated protein |
| TUBB | Tubulin beta class I |
| RUBCN | Rubicon autophagy regulator |
| RPS27A | Ribosomal protein S27a |
| ATG5 | Autophagy related 5 |
| SQSTM1 | Sequestosome 1 |
| CTTN | Cortactin |
| CTTNBP2 | Cortactin binding protein 2 |
| CTTNBP2NL | CTTNBP2 N-terminal like |
| ATG9A | Autophagy related 9A |
| MAP1LC3B | Microtubule associated protein 1 light chain 3 beta |
| ATG3 | Autophagy related 3 |
| GFAP | Glial fibrillary acidic protein |
| ALDH1L1 | Aldehyde dehydrogenase 1 family member L1 |
| SOX9 | SRY-box transcription factor 9 |
| GLUL | Glutamate-ammonia ligase |
| AQP4 | aquaporin 4 |
| KCNJ10 | Potassium inwardly rectifying channel subfamily J member 10 |
| GJA1 | Gap junction protein alpha 1 |
| GJA10 | Gap junction protein alpha 10 |
| NFIA | Nuclear factor I A |
| FABP7 | Fatty acid binding protein 7 |
| CD44 | CD44 molecule (IN blood group) |
| CCL2 | Chemokine (C-C motif) ligand 2 |
| HMGCS2 | 3-hydroxy-3-methylglutaryl-CoA synthase 2 |
| DIO2 | Iodothyronine deiodinase 2 |
| GLDC | Glycine decarboxylase |
| GPC5 | Glypican 5 |
